# Levels of DNA accessibility in human sperm vary across individuals with differing reproductive parameters

**DOI:** 10.1101/2024.06.27.600938

**Authors:** Mark E. Gill, Manuel Fischer, Christian De Geyter, Antoine H.F.M. Peters

## Abstract

Mammalian sperm DNA is packaged in a much denser form than in somatic cells, protecting the DNA from damage and reducing the size of the nucleus to improve passage towards the oocyte. While defective sperm DNA packaging correlates with reduced fertilization rates and impaired pre-implantation embryonic development, these defects are not currently examined in standard semen analysis. Here, we adapted NicE-view, an assay that directly labels accessible DNA, for use in human sperm and applied this method including extensive image quantification to examine spermatozoa from individuals with normal conventional semen parameters but variable reproductive outcomes. We found that two sub-populations of cells differing greatly in their DNA accessibility exist within both total and motile sperm. The frequencies of these two sub-populations vary between individuals, and selection of motile sperm by swim-up generally enriches for sperm with high DNA accessibility. Individuals with high frequencies of sperm with high DNA accessibility possess decreased sperm concentrations and increased DNA nicking levels and a subset of these individuals have a history of post-fertilization embryogenic failure. NicE-view shows much clearer separation of staining levels than alternative DNA labeling approaches, and represents a valuable tool for the assessment of human sperm.

## Introduction

Infertility is a widely prevalent condition and at least half of all cases being ascribed to male infertility^1^, though the accuracy of this characterization is unclear^2^. Aside from hormonal and anatomical sources, sperm-based defects are considered as the main causes. Current diagnostic criteria, established by the World Health Organization and based on epidemiological studies, consider sperm number, motility and morphology as standard parameters in the assessment of sperm function^3^. These features clearly associate with time-to-pregnancy (TTP), but fail to explain all infertility found in the population^4^. Additional assays examining other features of sperm have been proposed, but none have been globally accepted^3^.

One feature long associated with poor reproductive outcomes is sperm DNA fragmentation^5^. A recent meta-analysis has shown a link between increased DNA fragmentation in sperm and decreased clinical pregnancy rates^6^. However, the lack of standardization in how to measure this feature^7^ and the variation even within the use of a single assay^8^ has limited the global acceptance of DNA fragmentation as a standard diagnostic feature for male infertility.

An additional, and related, feature suggested to play a role in male infertility is sperm chromatin structure. Sperm chromatin is highly compacted when compared to somatic cells, a feature thought to both protect the DNA from exogenous damage and enable more efficient passage towards the oocyte. DNA in nearly all eukaryotic cells is packaged using highly conserved histone proteins, while mammalian sperm is mostly packaged with small, basic DNA binding proteins known as protamines^9^. Depletion of protamine is associated with increased sperm DNA damage and decreased fertility in both mice and humans^10–12^. Both fragmentation and reduced packaging of sperm DNA have been linked with developmental arrest of pre-implantation embryos^13–15^. A specific test of abnormal chromatin packaging in human spermatozoa would thus be extremely helpful in disentangling defects in the oocyte from those in the fertilizing sperm during early embryonic development.

Many studies of sperm chromatin utilize indirect assays to identify changes in DNA packaging proteins. For instance, increased levels of chromomycin A_3_ (CMA3), a DNA binding fluorescent dye, have been suggested to correlate with poor reproductive outcomes^16^. Staining by this dye in sperm has been associated decreased protamination^17^, though this connection is indirect at best. Bianchi et al. showed that CMA3 staining of fixed sperm from mouse and human could be quenched by pre-treatment of slides with salmon sperm protamine^17^. Other work has shown that zygotes generated by intracytoplasmic sperm injection (ICSI) from sperm enriched for strong CMA3 staining more frequently failed to decondense the sperm nucleus following fertilization compared to controls (where CMA3^high^ sperm was not abundant)^18^. Thus it is clear that the packaging of sperm chromatin is distinct and varies between species and individuals, but interpretation of the results is based on differences in correlation coefficients only.

Direct molecular assessment of chromatin accessibility is possible using a variety of approaches that enzymatically alter accessible DNA. These approaches include the <u>A</u>ssay for <u>T</u>ransposase <u>A</u>ccessible <u>C</u>hromatin (ATAC) methods (ATAC-Seq^19^ and ATAC-See^20^) where an engineered, hyperactive bacterial transposase (Tn5) integrates oligonucleotides into the DNA of accessible regions. ATAC-Seq has been performed in human^21^ and mouse^22^ sperm showing enrichment of accessible DNA regions near gene promoters and enhancers, though a recent study has questioned whether extracellular DNA contamination may influence these profiles in the mouse^23^. An alternative approach to label accessible DNA in intact cells are the so-called NicE techniques, where a nicking endonuclease generates single-stranded DNA nicks in accessible regions that then serve as initiation sites for *E. coli* DNA Polymerase I to synthesize down-stream stretches of DNA^24–26^. When biotinylated dNTPs are included with the polymerase, accessible DNA is labeled with biotin and can thus be captured by streptavidin and sequenced (NicE-Seq)^25^. NicE-Seq has been reported to generate results similar to those seen using ATAC-Seq, but with generally better resolution in fixed cells^25^. The NicE-view technique in contrast, where fluorescently labeled nucleotides are incorporated downstream of accessible, induced DNA nicks by DNA Polymerase I allows visualization of these sites in single cells^24,25^. To date no studies using NicE-view in sperm have been reported.

Here we assayed levels of accessible DNA using NicE-view in human sperm samples from individuals with different reproductive outcomes. Our aim was to use this assay to compare sperm from infertile individuals with similar fertilization rates but varying embryonic development rates upon application of ART. We found that NicE-view reveals variation in levels of accessible DNA in sperm from most samples, and that selection for highly motile sperm, via swim-up preparation, selects for cells with high levels of DNA accessibility in some individuals. Finally, we compared NicE-view to other assays for DNA accessibility and DNA damage and identified areas of overlap and difference in these assays.

## Materials and Methods

### Participant Selection

This study was approved by the Ethikkommission Nordwest-und Zentralschweiz (BASEC-ID 2020-00330). Study participants were recruited from individuals previously diagnosed and treated at RME (Universitätsspital Basel). Individuals were selected based on the presence of a previously obtained conventional semen analysis showing values within the WHO reference range^27^ (initial selection occurred prior to the publication of the sixth edition sperm analysis manual in 2021^3^). Partners of selected individuals had normal antral follicle counts and AMH values within standard range. Individuals in the naturally occurring pregnancy (NATP) group obtained a pregnancy following an initial semen analysis either spontaneously or following a non-ART treatment modality. Individuals in the high blastocyst growth rate (HBGR) and low blastocyst growth rate (LBGR) groups were selected from those who underwent ART-based treatment at RME (either ICSI or IVF) between 2018 and 2021. In all individuals selected the rate of generation of zygotes following ART was normal. Inclusion in the HBGR group was indicated by the production of blastocysts from at least 50% of cultured embryos, while LBGR group inclusion was defined by generation of at most 1 blastocyst following embryo culture.

Samples used for FACS isolation of sperm based on Chromomycin A_3_ (CMA3) levels were obtained from participants from an additional cohort (recruitment of which was also approved by the Ethikkommission Nordwest-und Zentralschweiz (EKNZ 2017-01407)) and whose sperm were previously examined for alterations in DNA methylation^28^. This cohort consisted of a control group of fertile individuals who had previously donated sperm (and whose semen parameters were within the WHO standard ranges) and a group of infertile individuals with normal semen parameters, but anogenital distance (AGD) of less than 40 mm^29^ and >20% TUNEL positive swim-up sperm in a previous donation.

All participants provided informed consent prior to providing semen samples for analysis.

### Participant data analysis and statistics

Participant data was extracted from a clinic database and coded to remove all personally identifiable information with coding only available to clinical personnel (C.D.G.). Coded data were then entered into a secuTrial database (interActive Systems GmbH, Berlin Germany), which was accessible to researchers. Data was extracted from this database and analyzed using R^30^. Summary tables and statistical calculations were generated using the gt^31^ and gtsummary^32^ R packages.

### Sperm sample collection and processing

Participants were requested to abstain from sexual activity for at least 2 days prior to providing a semen sample via masturbation at RME (Universitätsspital Basel) (actual abstinence times ranged from 1-14 days). Semen samples were allowed to liquefy at 37°C and then processed for conventional semen analysis in the Andrology laboratory at RME^27^. A portion of each semen sample was subjected to a swim-up preparation to enrich for motile sperm^33^. In 57 of 60 cases sufficient motile sperm was obtained for analysis, while swim-up preparation from 3 individuals resulted in insufficient material for further experimentation. For this study total sperm refers to liquefied semen that was washed once in Ham’s F-10 (MHF-10) media. A fraction of total sperm was used for a TUNEL assay for DNA fragmentation^8^. Total and swim-up sperm were dried onto 10-well diagnostic slides (VWR, Radnor USA, #631-1371; 30,000 sperm per well) and then transferred to -20°C for storage.

### NicE-view staining of human sperm samples

10-well slides of total and swim-up sperm were thawed at room temperature and washed once with phosphate buffered saline (PBS, 50 µl per well). Sperm were then fixed with 1% paraformaldehyde (Electron Microscopy Sciences, Hatfield USA, #15713) in PBS for 10 minutes at room temperature, followed by two washes in PBS. Sperm were permeabilized using Omni ATAC permeabilization buffer^34^ (10 mM Tris, pH 7.5, 10 mM NaCl, 3 mM MgCl_2_, 0.1% Igepal, 0.1% Tween-20, 0.01% digitonin) without DTT for 15 minutes at room temperature and then washed once with PBS. For T4 NicE-view, following permeabilization samples were incubated for 15 minutes at room temperature with 30 µl per well of T4 DNA ligase (ThermoFisher Scientific, Waltham USA, #EL0012) in 1x T4 DNA ligase buffer (40 mM Tris-HCl, 10 mM MgCl_2_, 10 mM DTT, 0.5 mM ATP, pH 7.8) and subequently washed once with 50 µl per well PBS. Samples were then labeled for 2 hours at 37°C in 30 µl per well labeling mix. The NicE-view labeling mix was based on previously published protocols^24–26^ and consisted of 1x NEBuffer 2 (NEB, Ipswich USA, #B7002S), 30 µM dGTP (ThermoFisher Scientific, #R0181), 30 µM dTTP (ThermoFisher Scientific, #R0181) and 30 µM dmCTP (Jena Bioscience, Jena Germany, #NU-1125L), 24 µM dATP (ThermoFisher Scientific, #R0181), 6 µM Fluorescein-dATP (Jena Biosciences, #NU-1611-FAMX), 3.2 U/ml Nt.CviPII (NEB #R0626S) and 64 U/ml *E. coli* DNA Polymerase I (NEB #M0209L). For Nick translation the nicking endonuclease Nt.CviPII was excluded, while for negative control reactions both Nt.CviPII and DNA Polymerase I were excluded. Following labeling, samples were washed 3 times with PBS + 0.01% SDS + 50 mM EDTA, pre-heated to 55°C, for 15 minutes each. Samples were counter-stained with DAPI (10 µg/ml in PBS, Sigma Aldrich, St. Louis USA, #D9542-10MG) for 5 minutes at room temperature, mounted with VectaShield (Vector Laboratories, Newark USA, #H-1000) and covered with 50 mm coverslips. Slides were then sealed with nail polish. Slides were stored at 4°C until imaging (maximally 2 days).

### Double labeling of DNA nicks & NicE-view signal on human sperm

For double labeling, sperm were processed as for standard NicE-view labeling with the following exceptions. Following fixation and permeabilization, endogenous DNA nicks were labeled using Fluorescein-labeled dATP (as above). Ends of newly synthesized DNA strands were then sealed by incubating with T4 DNA ligase (ThermoFisher Scientific, EL0012) in 1x T4 DNA ligase buffer overnight at room temperature in a humid chamber. A second labeling reaction with ATTO565-labeled dATP (Jena Biosciences, #NU-1611-565) was then performed either with Nt.CviPII or without (as control). Post-labeling processing was as in standard NicE-view labeling (described above).

### FACS sorting of sperm based on CMA3 levels and microscopic analysis

Total sperm from fertile participants and infertile participants with decreased anogenital distance and increased percentages of TUNEL positive was subjected to sorting based on CMA3 levels as previously described^28^. Briefly, total, media-washed sperm was incubated in the dark for 30 minutes at room temperature with 0.25 mg/ml Chromomycin A_3_ (CMA3, Sigma Aldrich, #C2659-10MG) and 5 µM Vybrant DyeCycle Ruby (ThermoFisher Scientific, #V10273) diluted in McIlvaine’s buffer, pH 7.0, supplemented with 10 mM MgCl_2_. Sperm were then washed once with diluent.

FACS was performed using a BD Influx (BD Biosciences, Franklin Lakes USA) equipped with 355 nm, 488 nm and 640 nm lasers. Cells were first gated using FSC and SSC to eliminate debris. Haploid sperm were then further selected from these selected cells on the basis of DNA content (using Vybrant DyeCycle Ruby). Finally the selected sperm were sub-divided into two populations based on CMA3 levels. These selected sperm were then dried onto 10-well diagnostic slides and stored at - 20°C. Slides were later subjected to T4 NicE-view as described above.

### ATAC-See staining of human sperm

Oligonucleotides for ATAC labeling were synthesized and HPLC purified by Microsynth AG (Balgach Switzerland). The two forward oligos (TCGTCGGCAGCGTCAGATGTGTATAAGAGACAG & GTCTCGTGGGCTCGGAGATGTGTATAAGAGACAG) were labeled at their 5’ ends with ATTO488 (ATTO-Tec Siegen Germany), while a common reverse oligo (CTGTCTCTTATACACATCT) was 5’ phosphorylated. Forward and reverse oligos were annealed at a concentration of 1.5 µM in annealing buffer (10mM Tris pH 7.5, 50 mM NaCl and 1 mM EDTA) by heating to 95°C for 5 minutes and subsequently cooling to 25°C over a period of 45 minutes in a thermocycler.

Recombinant ZZ-tagged Tn5^35^ was purified from *E. coli*. Tn5 (∼5 µM) was loaded with labeled annealed oligos (6 µM) by mixing and incubating at 37°C for 30 minutes. Equimolar volumes of Tn5 loaded with each of the two different annealed oligos was pooled and stored at -20°C until use.

Pre-processing of 10-well slides was the same as for NicE-view (up to permeabilization). Transposition reactions were performed in 1x TDM buffer (10 mM Tris, pH 7.5, 10% DMF, 5 mM MgCl_2_). Fluorescent oligo-loaded Tn5 was diluted 1:100 for use. For the no Magnesium control 5 mM MgCl_2_ was replaced by 5 mM EDTA. Transposition reactions were performed at 37°C for 1 hour in a humid chamber. Post-staining processing (washing, counter-staining and mounting) was the same as for NicE-view.

### NicE-view and ATAC-See imaging

Stained slides were imaged on a Visitron Spinning Disk W1 confocal microscope (Visitron Systems GmbH, Puchheim Germany). Images were acquired with a 40x oil objective (NA: 1.3, WD 0.21) using a 405 nm laser with a 460/50 filter set for DAPI and a 488 nm laser with a 525/50 filter set for NicE-view and ATAC signal. For double labeling experiments a 561 nm laser with a 609/54 filter set was also utilized. For each well, 2 separate 5x5 tiles were acquired including a 6 µm Z-stack for each image (with 1 µm step size). For Nick translation, NicE-view and T4 NicE-view, 2-wells per slide were independently stained resulting in 100 image stacks per slide for these assays. For negative control and T4 DNA ligase treatment of Nick translation, 1 well per slide was stained resulting in 50 image stacks per slide for these controls.

### NicE-view and ATAC-See image processing

Maximum intensity projections were generated for each stack in VisiView (Visitron Systems, Puchheim Germany). Projections were processed with Fiji^36^ and Photoshop (Adobe Systems, San Jose, USA) to generate images shown in Figure 1B and Supplementary Figure S1B. Projections were further processed using CellProfiler^37,38^. Nuclear segmentation was performed on DAPI images (Segmentation Parameters: 25-50 pixel diameter, Adaptive Sauvola thresholding with no smoothing scale, a threshold correction factor of 1.0, and an adaptive window size of 800. Clumped objects were distinguished by shape with intensity used to draw dividing lines between clumped objects). Segmentation masks were inspected visually to confirm reasonable segmentation. Intensity values and size and shape parameters were then calculated for each segmented nucleus.

**Figure 1.**
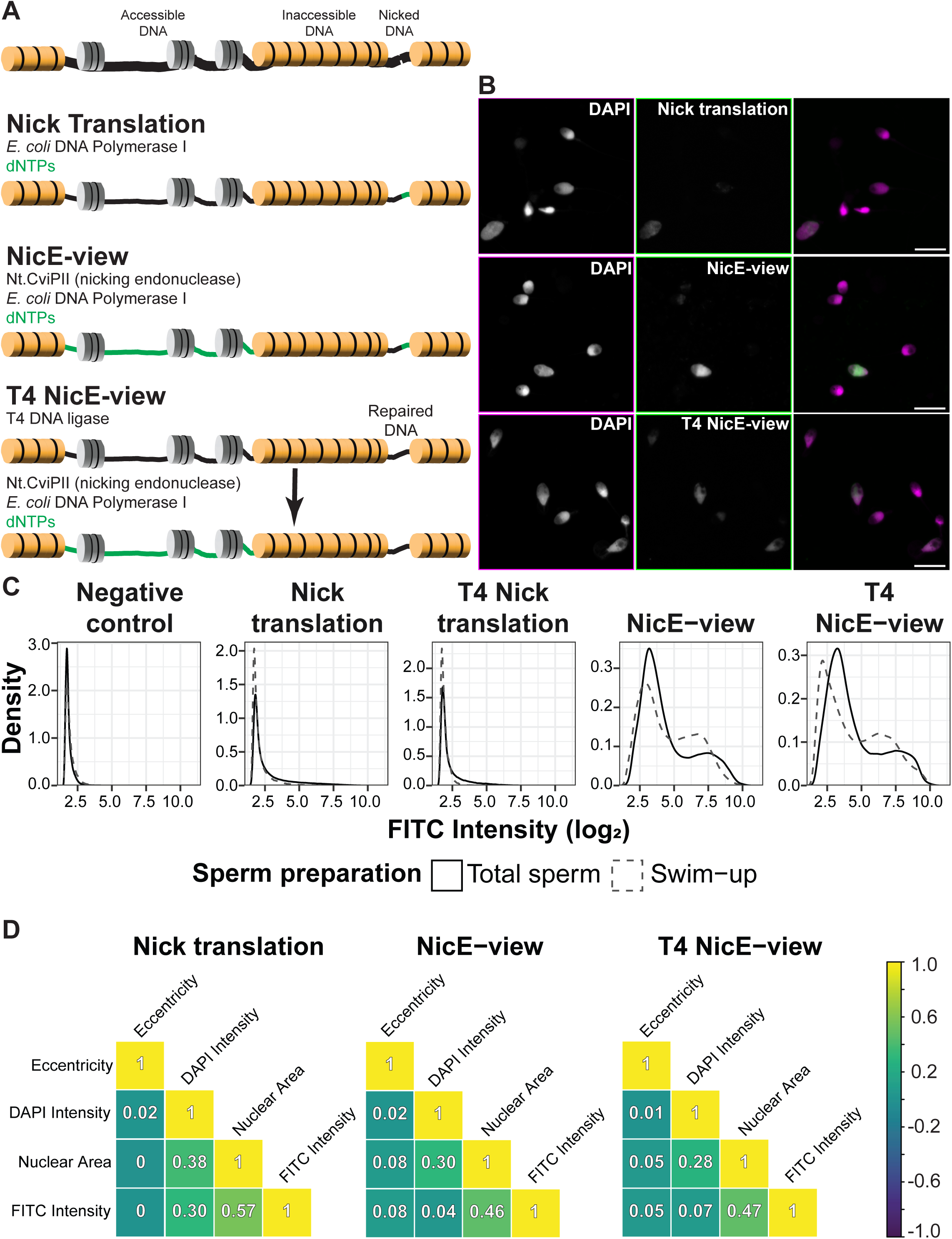
NicE-view reveals variation between DNA accessibility in individual human sperm. (A) Cartoon showing different staining conditions used to measure presence of accessible DNA and nicked DNA in human sperm samples. (B) Example images showing results of staining outlined in (A). Brightness and contrast for Nick translation image were enhanced relative to others to make signal more visible (Supplementary Figure S1 shows unenhanced image). Scale bars = 10 μm. (C) Density plot showing distributions of log2 transformed integrated intensities for sperm stained with conditions indicated in (A). (D) Correlation plots showing

### Imaging data analysis

All data analyses were performed in R^30^. Comma separated value (csv) files generated by CellProfiler were loaded into R and pre-processed using the tidyverse suite of packages^39^.

Correlation matrices and their significance values were generated using the rstatix package (cor_mat parameters: method = “spearman”, alternative = “two.sided”, adjust_pvalue(method = “fdr”)40.

Thresholding of Nick translation, NicE-view, and T4 NicE-view distributions was performed as follows. A kernel density estimation was performed for log_2_- transformed signal intensity values for each sample and staining condition (in a range of median ±2). Optimization was performed on an approximated function derived from this estimation and the minimum value from this optimization was determined. For NicE-view and T4 NicE-view this value was used as the threshold to separate sperm into highly or lowly stained categories. For Nick translation, the value calculated for the Negative control of the same sample was used as the threshold value.

### Data visualization

Most plots were generated using ggplot2^41^. The ggpubr package was used to add correlation coefficients to scatter plots^42^. Beeswarm plots (Figure 2B) were generated using the ggbeeswarm package^43^. Plots were combined using the patchwork package^44^. For plots showing distributions of values between individuals, 1000 sperm were randomly sub-sampled (using sample_n()^39^) from each sperm sample (total and swim-up) to generate distributions equally weighted for each individual. Scatter plot shown in Figure 5D was generated using the scattermore package^45^.

**Figure 2.**
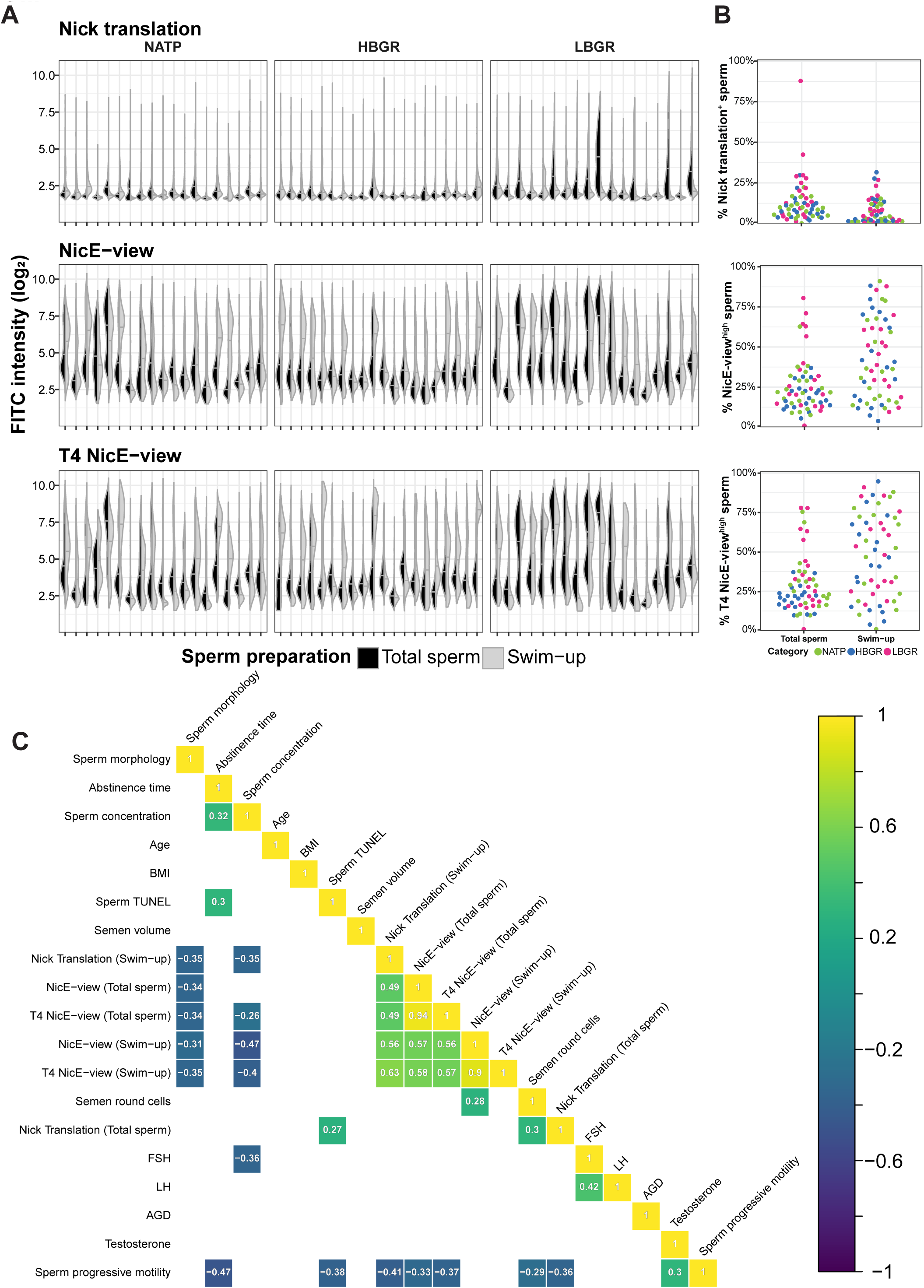
Frequency of sperm with differing levels of DNA accessibility vary between individuals and preparation methods. (A) Split violin plot showing the distribution of single sperm integrated intensity values from 57 individuals stained with Nick translation, NicE-view and T4 NicE-view. Black distributions (left) represent total sperm, while grey distributions (right) refer to sperm selected for motility via swim-up preparation. (B) Beeswarm plot showing distributions of frequencies of highly labelled sperm (for each assay) obtained from thresholding distributions seen in (A), colored by participant category. For total sperm n = 60, for swim-up n = 57. (C) Correlation plot showing Spearman’s ρ for a set of participant level features. Values are

Correlation plots (Figure 1D, Supplementary Figure 1C & Figure 2C) were generated using the cor_plot() function from the rstatix package^40^. Colors were generated using the viridis package^46^. For Figure 2C only values with an FDR-corrected p < 0.05 were displayed.

Table (Figure 3A) showing statistical significance of parameters upon re-categorization by NicE-view behavior was generated using the gtsummary package32.

**Figure 3.**
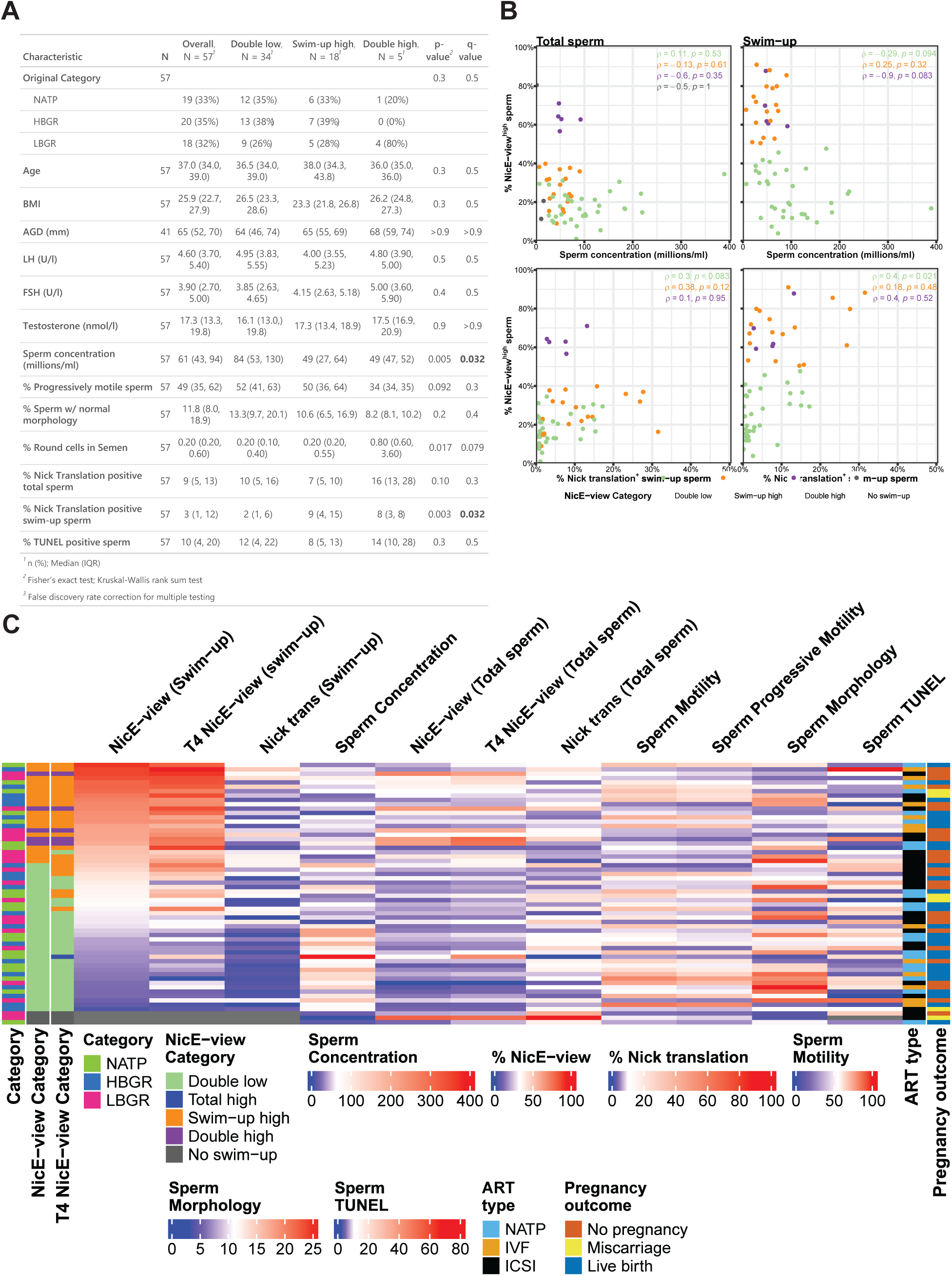
Sperm concentration is negatively correlated with the frequency of sperm with high levels of DNA accessibility. (A) Table showing breakdown of participant parameters following re-categorization based on NicE-view frequencies (from Figure 2B). Data from 57 individuals possessing both total and swim-up samples is shown. Double low: <50% of sperm with high DNA accessibility in both total and swim-up sample. Double high: ≥50% of sperm with high accessibility in both total and swim-up sample. Swim-up high: ≥50% of sperm with high accessibility In swim-up sample, but not in total sample. For numerical variables, a Krus-kal-Wallis rank sum test was used to compare groups, while for Original Category a Fisher’s exact test was used. Statistical comparisons of different parameters were corrected for multiple hypothesis testing using a False discovery rate calculation (listed as “qvalue” in table). (B) Scatter plots comparing Sperm concentration or % Nick translation^+^ swim-up sperm to % NicE-view^high^ sperm (from total (left) and swim-up (right)). Points are colored based on NicE-view Categ ries as defined in (A). Values shown are Spearman’s ρ by category. (C) Heatmap showing relationship between spermatological parameters, DNA accessibility and DNA nicking in all

Heatmaps (Figure 3C, Supplementary Figure S2A) were generated by the ComplexHeatmap package^47,48^. Color scales for each numerical variable were generated by the colorRamp2() function from the circlize package^49^ such that median values were colored white, while minimum values were blue and maximum values red. Row order for heatmaps were determined by the frequency of NicE-view^high^ swim-up sperm.

### Data availability

Quantitation of imaging data and code used for analysis are available at Zenodo (10.5281/zenodo.12526509).

## Results

### Selection of individuals with sperm of differing embryonic competence

To study the role of sperm-based factors in post-fertilization embryonic development we recruited a group of individuals previously diagnosed and treated for infertility in our clinic. Selected individuals were chosen based on values of conventional semen analysis obtained at the time of diagnosis (labeled as “historical sample” in Table 1) that were largely within the reference range of spermatology values established by the WHO^27^.

**Table 1.**
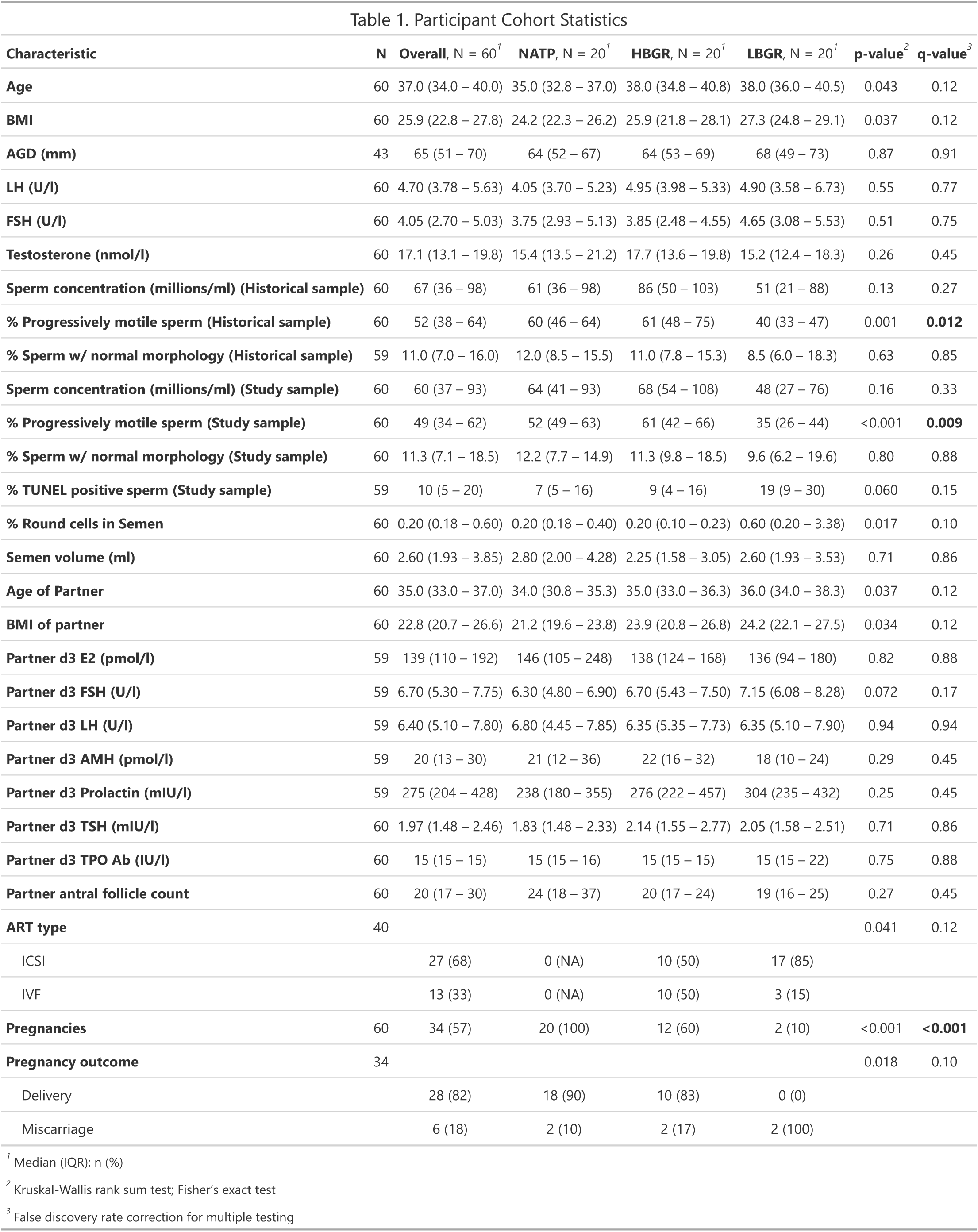
Characterization of participant cohort. Demographic, hormonal and reproductive parameters for participants recruited for this study. Historical sample refers to sperm sample obtained at time of initial diagnosis, while study sample refers to sample obtained as part of this study (and subsequently used for further analyses). Partner parameters refer to values obtained from oocyte providing partners of study participants. For numerical variables, a Kruskal-Wallis rank sum test was used to compare groups, while for categorical variables (ART type, Pregnancies and Pregnancy outcome) a Fisher’s exact test was used. Statistical comparisons of different parameters were corrected for multiple hypothesis testing using a False discovery rate calculation (listed as “q-value” in table).

One group of 20 individuals initially sought treatment, but obtained a natural pregnancy without the use of assisted reproductive technologies (ART) (referred to as NATP). Two additional groups of 20 individuals each were treated with ART and achieved normal fertilization rates, but showed variation in pre-implantation embryonic development. For one group the development of zygotes to the blastocyst stage was substantial (>50%, referred to as high blastocyst growth rate or HBGR). The other ART-treated group consisted of couples in which the development of zygotes to blastocyst was severely impaired (maximally 1 blastocyst formed, referred to as low blastocyst growth rate or LBGR).

Comparison of features between participant groups showed no significant differences in age, BMI or anogenital distance (AGD) (Table 1). Hormonal parameters (LH, FSH and testosterone) were also not significantly different between groups (Table 1).

Comparison of semen parameters between groups showed a significant decrease in the percent of progressively motile sperm in the LBGR group relative to the two others, but no differences in any other parameters (Table 1). Despite this, the sperm samples (acquired at time of recruitment) of 15 of 20 LBGR individuals possessed progressive motility values above the 5^th^ percentile reference values for fertile individuals as determined by the WHO^27^.

Analysis of age, BMI, endocrine features and antral follicle counts in partners of study participants showed no significant differences between groups, minimizing the possibility for ovarian reserve or oocyte-derived factors in driving differences in pre-implantation development rates in these couples (Table 1).

### NicE-view reveals two primary sub-populations of sperm in human samples

To examine the chromatin state of sperm we modified the NicE-view assay^24–26^ to label accessible DNA in frozen, dried sperm samples. This assay introduces single-stranded DNA nicks into regions of accessible chromatin and then labels areas surrounding these nicks via introduction of labeled nucleotides by *E. coli* DNA polymerase I^24–26^ (Figure 1A). We first optimized permeabilization of the sperm, using a modified version of the permeabilization buffer developed for Omni ATAC-Seq, which has been shown to work in a variety of fixed cells^34^. For use in sperm we removed DTT from this buffer, to prevent reduction of disulfide bonds in protamines and possible alteration of chromatin structure in these cells. NicE-view cannot distinguish between endogenous single-stranded DNA nicks and accessible chromatin. It has previously been reported that some human sperm contains substantial levels of endogenous DNA nicks^17,50^, a result we confirmed by examining DNA nicking using Nick translation (Figure 1A&B). We modified NicE-view to more specifically label only accessible chromatin by treating samples with T4 DNA ligase prior to NicE-view labeling (referred to as T4 NicE-view). We evaluated whether this pre-treatment was effective at repairing DNA nicks in this context by treating samples with T4 DNA ligase prior to performing a Nick translation assay (referred to as T4 Nick translation, Supplementary Figure S1A&B). We found that this pre-treatment was indeed effective at supressing signal from Nick translation, though this supression was less efficient in samples with very high signal, where DNA is likely more highly fragmented (Figure 1C). In brief: Nick translation reflects the quantity of endogenous DNA nicks, NicE-view reflects accessible portions of the genome and regions possessing endogenous DNA nicks and T4 NicE-view reflects only accessible portions of the genome.

Imaging of human sperm with the modified NicE-view protocols (with and without T4 DNA ligase) revealed a wide variation in signal intensities between individual spermatozoa within each sample (Figure 1A). Generally, signal intensity for the NicE-view (and T4 NicE-view) labeled samples was much higher than that seen with Nick translation (Figure S1B, note settings for unenhanced image is identical to NicE-view and T4 NicE-view in Figure 1B). We imaged sperm labeled via Nick translation, NicE-view and T4 NicE-view (and controls) from 117 samples (derived from 60 individuals: 60 total washed semen samples (referred to as total sperm) and 57 sperm samples prepared using the swim-up method to select for motile sperm^33^) by confocal microscopy.

To obtain a quantitative view of DNA labeling in individual sperm, we performed segmentation and quantification of nuclear signal from these images leading to measurements of 3,376,931 individual spermatozoa. Plotting the density of signal intensities across different staining conditions showed that negative controls (treated with fluorescently labeled dNTPs but no enyzmes) possessed very low background signals, while Nick translation signal was similar in intensity to background levels for the majority of sperm, with only a small population showing a relatively high signal (Figure 1C). Sperm with a high Nick translation signal were more abundant in total sperm when compared to swim-up prepared sperm (Figure 1C), consistent with previous studies showing that swim-up selection decreases the frequency of DNA fragmentation in sperm^51,52^. T4 DNA ligase pretreatment did not fully suppress the high Nick translation signal in brightly labeled sperm, a notion more clearly seen in total sperm (Figure 1C), suggesting that within the total sperm population a group of sperm with high levels of DNA nicks is present and mostly removed via selection for sperm motility.

In contrast to Nick translation, where the majority of sperm showed signal near to background, the vast majority of sperm stained via NicE-view and T4 NicE-view showed clear labeling (Figure 1C). We observed a bimodal distribution of signal intensities in both of these assays, an effect present in both total and swim-up sperm (Figure 1C). We also observed that the values for the two peaks of signal intensities for swim-up sperm were slightly decreased relative to total sperm in both assays, though the frequency of sperm in the upper portion of the distribution was increased in swim-up prepared samples (Figure 1C).

We also quantified a variety of features related to DNA labeling, DAPI staining intensity and the size/shape of the nuclei of the spermatozoa. Correlation analysis of many of these features showed strong correlations as they are highly interdependent variables (Supplementary Figure S1C). We selected three measurements in addition to integrated DNA labeling intensity (which sums the value across the entire nucleus, labeled as FITC intensity in Figure 1D) representing basic features of sperm nuclei: (1) Nuclear area, (2) Eccentricity (representing the level of nuclear elongation), and (3) integrated DAPI intensity (representing total DNA staining levels). Analysis of these four variables showed a clear correlation between Nuclear area and DNA staining intensity (regardless of staining modality) (Figure 1D). This effect may be expected for integrated intensity measurements, as summing even background intensity values across more pixels would lead to increased signal, however, we also observed a correlation between mean staining intensity and nuclear area (Supplementary Figure S1C), which should not be driven by the increase in pixel number. We also observed correlations between the extent of enzymatic DNA labeling (by FITC incorporation) and nuclear area for all 3 staining approaches (Figure 1D). DNA dye staining and the extent of DNA labeling showed a variable effect based on staining condition: while signal intensity derived from endogenous nicks showed a correlation with total DNA staining level, accessible DNA labeling showed no such effect (Figure 1D). None of the selected variables showed a correlation with nuclear shape as measured by eccentricity (Figure 1D).

### DNA Nicking and NicE-view labeling vary between individuals and correlate with reproductive parameters

We next asked how the distribution of NicE-view signal intensities varied between individual study participants. This analysis showed a wide variety of values across participants (Figure 2A). We observed a few patients of the LBGR sub-group with relatively high levels of endogenous DNA nicking in their total sperm (which were largely absent following selection via swim-up) (Figure 2A). We also observed a tendency towards higher frequencies of sperm with increased DNA accessibility (measured both with NicE-view and T4 NicE-view) in swim-up samples from all cohort sub-groups (Figure 2A). In addition, there was a tendency towards increased frequencies of highly accessible DNA in total spermatozoa of individuals from the LBGR sub-group compared to the others (Figure 2A).

To compare values from our labeling assays to other reproductive parameters, we utilized a thresholding approach to sub-divide sperm into groups based on signal intensity. To threshold Nick translation signal we identified a non-zero local minima in our Negative Control distributions for each sample and applied these as cutoff values defining sperm with greater than these values as Nick translation positive. For NicE-view and T4 NicE-view we identified the local minimum around the median signal intensity value for each distribution in each sample. We defined sperm with values above the local minima as NicE-view^high^ and those with values below as NicE-view^low^.

This thresholding clarified the trends shown in Figure 2A. The number of total sperm with DNA nicks was clearly higher than that seen in swim-up selected sperm, with a subset of LBGR individuals showing higher values in total sperm relative to the other participants (Figure 2B).

The distribution of NicE-view^high^ (and T4 NicE-view^high^) sperm frequencies in total sperm show that only ∼10% samples possessed greater than 50% NicE-view^high^ sperm (Figure 2B). Among the individuals with greater than 50% NicE-view^high^ total sperm 5/6 were LBGR and for T4 NicE-view this was 6/7 (Figure 2B). In contrast, following swim-up selection ∼40% of individuals have frequencies of NicE-view^high^ sperm greater than 50% (Figure 2B). No participant category was found to have an enrichment in either the high or low frequency groups for swim-up samples (Figure 2B).

We next determined the correlation between the frequencies of DNA-nicked or DNA-accessible sperm and other reproductive parameters (Figure 2C). We found strong correlations between NicE-view^high^ and T4 NicE-view^high^ frequencies in both total and swim-up sperm (Figure 2C).

We also found that the frequency of swim-up sperm with DNA nicks, but not total sperm, positively correlated with DNA accessibility (Figure 2C). The frequency of Nick translation-positive total sperm showed a positive correlation with an independent measure of DNA fragmentation (TUNEL, Terminal dUTP Nick End labeling, which measures both single and double-stranded DNA damage^8^), but this correlation disappeared in swim-up selected sperm (Figure 2C).

Finally, we found that sperm parameters measured as part of conventional semen analysis are negatively correlated with the frequency of high DNA accessibility sperm across all samples (Figure 2C).

### Re-categorization of participants based on DNA accessibility

Our measurements of high DNA accessibility frequencies in total and swim-up sperm suggested a bimodal distribution of values (Figure 2B). Using a frequency of 50% as a cutoff, we re-categorized our participants based on the behavior of their sperm in both preparation methods. Using NicE-view^high^ frequencies, 3 categories can be defined: Double low (where <50% of sperm in both total and swim-up samples were NicE-view^high^), Double high (where ≥50% of sperm in both total and swim-up samples were NicE-view^high^) and Swim-up high (where ≥50% of swim-up sperm but <50% of total sperm were NicE-view^high^). 59.6% of individuals were categorized as Double low, 8.8% of individuals as Double high and 31.6% of individuals as Swim-up high (Figure 3A).

Comparing our newly defined categories to those used for recruitment, we found that 80% of NicE-view Double high participants were LBGR, however, this enrichment is not statistically significant owing to the low number of Double high individuals identified (Figure 3A). Comparing our NicE-view-defined categories to other participant parameters, we found two which were significantly different between categories: Sperm concentration and % Nick translation positive swim-up sperm (Figure 3A).

Comparing NicE-view^high^ frequency values to sperm concentration for each individual showed that no individual from the Double high or Swim-up high categories had a sperm concentration of greater than 100 million/ml, while 38% of individuals from the Double low category possessed a value above this level (Figure 3B). Comparison of swim-up Nick translation frequencies to NicE-view frequencies showed that while 20% of Double low individuals had greater than 10% Nick translation positive swim-up sperm, 50% of Swim-up high individuals had swim-up Nick translation frequencies above this value (Figure 3B).

Finally to compare all parameters and categories to each other, we plotted these data as a heatmap (Figure 3C). The relationships between NicE-view, T4 NicE-view and Nick translation values from swim-up sperm were clear (particularly for those with low frequencies for all parameters) (Figure 3C). The negative relationship between Swim-up NicE-view^high^ frequency and Sperm concentration was also clear (Figure 3C). We did not observe an obvious relationship between any other measured parameters (Figure 3C, Supplementary Figure S2A).

### Relationship between NicE-view and other methods for detecting DNA accessibility

Previous studies have found variation in the staining level of the DNA dye Chromomycin A_3_ (CMA3) between sperm from the same sample^17^. This variation has been suggested to represent variation in protamine levels between sperm^17^ and has been correlated with poor reproductive outcomes^16^. To determine if NicE-view^high^ sperm and CMA3^high^ sperm represent the same sub-population, we used FACS (Fluoresecence Activated Cell Sorting) of sperm stained with CMA3 to obtain CMA3^low^ and CMA3^high^ enriched sperm populations^28^. We then performed our T4 NicE-view assay on these two populations (isolated from 6 individuals). We found a general positive correlation between the two assays (Figure 4A; 5 of 6 individuals show increased median T4 NicE-view signal intensities in CMA3^high^ compared to CMA3^low^ sub-populations). However, the presence of clear bimodal T4 NicE-view intensity distributions in both CMA3 sub-populations suggests that these two sub-populations do not fully overlap (Figure 4).

**Figure 4.**
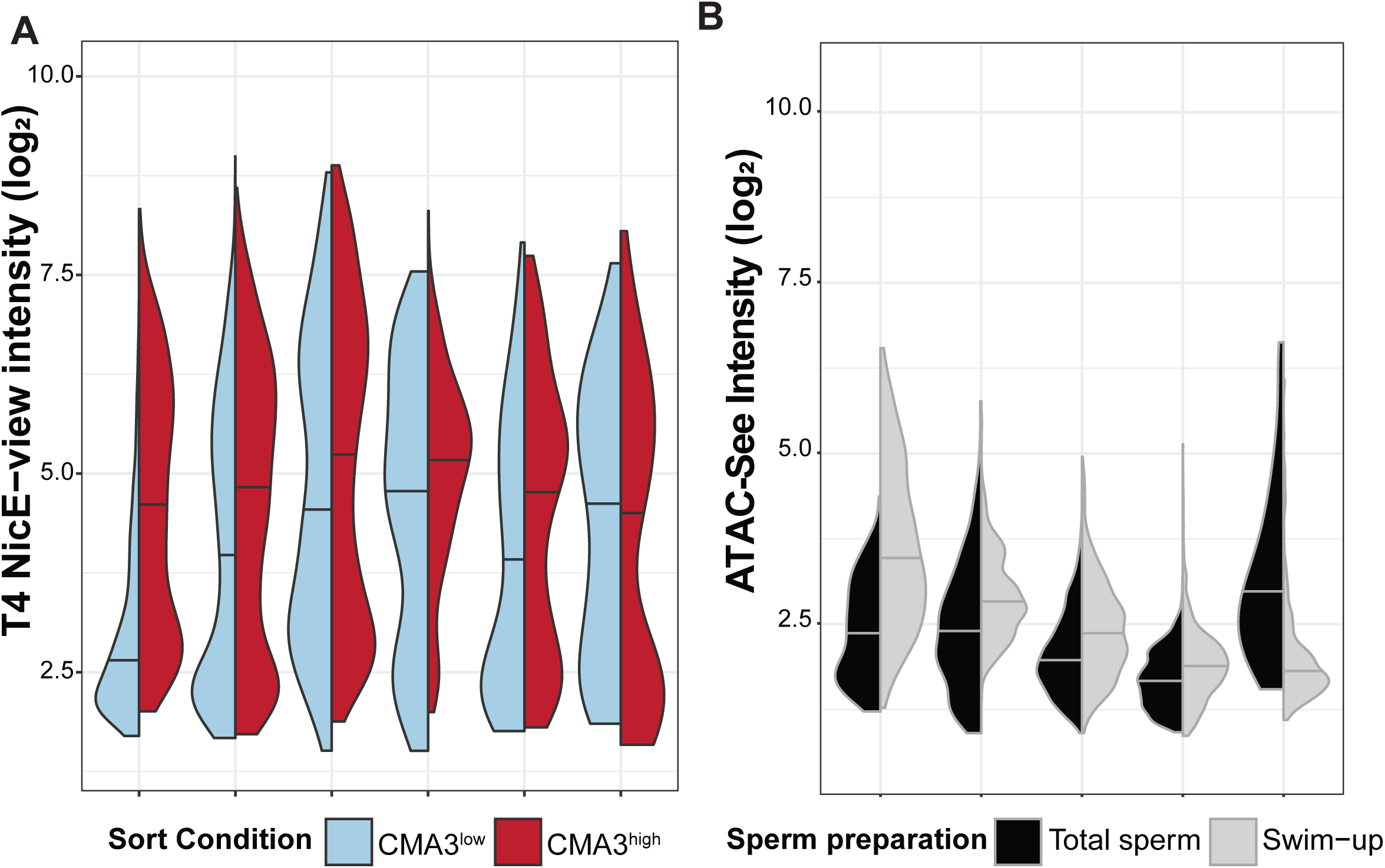
NicE-view intensity partially correlates with CMA3 and provides better separation of lowly and highly labelled sperm than ATAC-See. (A) Split violin plot showing the distribution of log2 integrated T4 NicE-view intensities from individual sperm from samples isolated by FACS for CMA3 from six individuals. Sperm with low CMA3 levels are shown in blue on left, while sperm with high CMA3 levels are shown in red on right. (B) Split violin plot showing the distribution of log2 integrated ATAC-See intensities from individual sperm from total and swim-up preparations from five individuals.

ATAC has been used extensively to measure DNA accessibility in many cell types including sperm^19–22,34^. We utilized ATAC-See^20^ (a fluorescence microscopy-based variation of the ATAC labeling protocol) to measure DNA accessibility in human sperm. We first tested whether incorporation of fluorescently labeled oligonucleotides into chromatin could occur in fixed sperm samples. We performed two control reactions to test for incorporation: first we excluded Tn5 transposase (which showed that amount of fluorescence exclusively obtained from labeled oligonucleotdies) and second we included Tn5, but excluded Magnesium (which is a necessary cofactor for Tn5’s transposition activity). Comparing these controls to ATAC-See reactions, we observed an increased signal intensity in ATAC-See showing that incorporation of oligonucleotides did occur in fixed human sperm samples, though the level of signal enrichment compared to the negative controls was relatively mild (Supplementary Figure S3A).

When we examined the levels of ATAC-See signal in total and swim-up sperm from 5 individuals we observed that 4/5 individuals showed increased median signal in swim-up compared to total sperm (Figure 4B). However, in contrast to NicE-view and T4 NicE-view we did not observe a bimodal distribution of signals and overall signal intensities were quite low (Figure 4B).

### Some, but not all, sperm with high DNA accessibility contain DNA nicks

Given the positive correlation between NicE-view^high^ and swim-up Nick translation frequencies (Figures 2C&3A), we next asked whether this correlation was driven by a common sub-population of sperm. Examination of individual samples showed a very strong correlation between frequencies of both NicE-view^high^ and T4 NicE-view^high^ sperm in both total and swim-up samples (Figure 5A). In contrast, no correlation between the frequency of NicE-view^high^ sperm and Nick translation positive sperm was found in total sperm, while a positive correlation was found for NATP and HBGR individuals in swim-up samples (Figure 5B).

**Figure 5.**
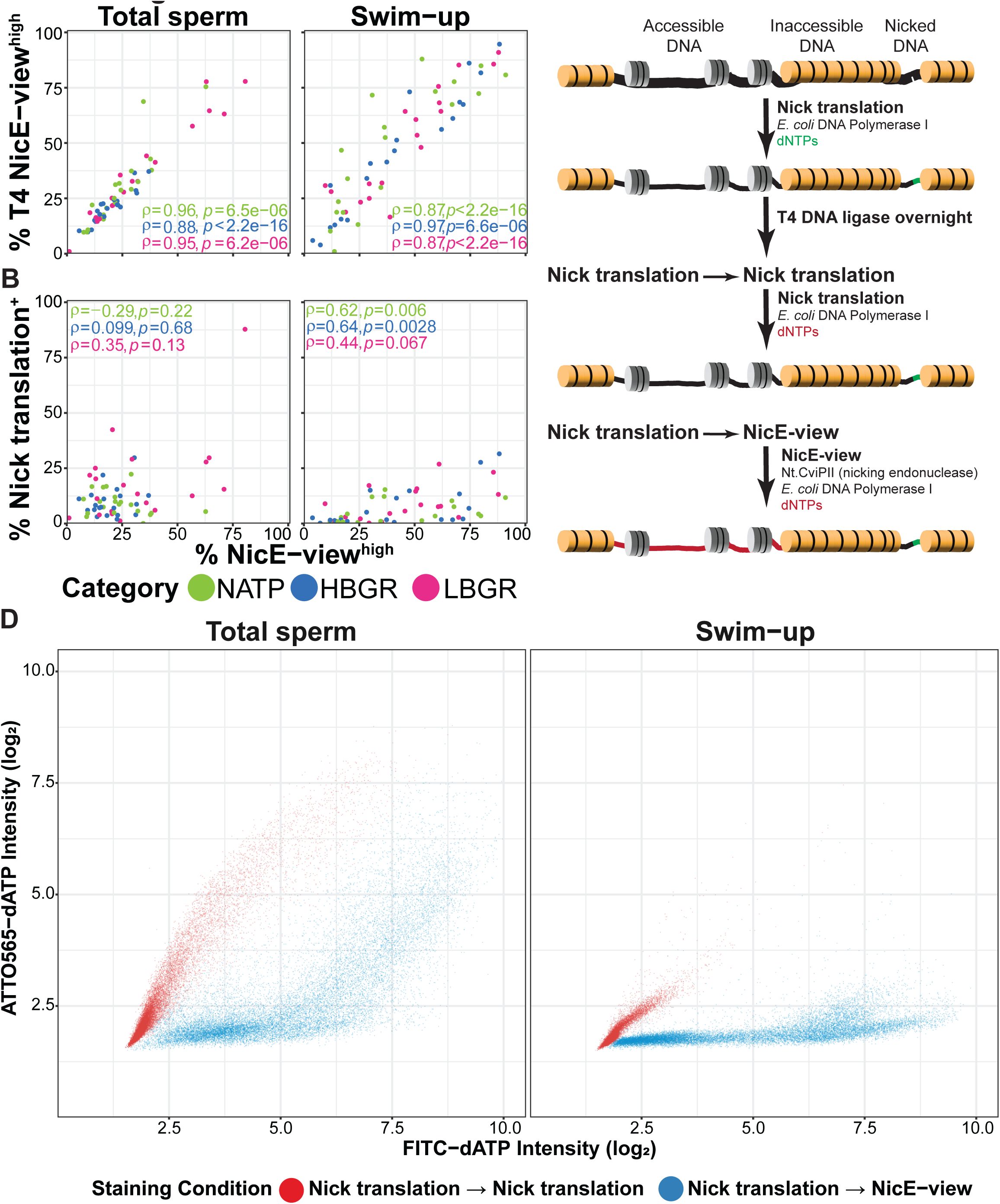
DNA accessibility and DNA nicking correlate in human sperm samples. (A&B) Scatter plot showing relationship between % NicE-view^high^ and % T4 NicE-view^high^ (A) or % Nick translation^+^ (B) sperm in total (left) and swim-up (right) samples. Points are colored by participant category. Values presented are Spearman’s ρ per category. (C) Cartoon describing double labeling experiment to label DNA nicks and accessible regions in the same cell. (D) Scatter plot showing the results of double labeling experiments. Red points indicate Nick translation followed by Nick translation, while blue points indicate Nick translation followed by NicE-view. For both samples value on y-axis represents Nick translation level. For red points value on x-axis represents incomplete DNA repair by T4 DNA ligase (likely from highly damaged sperm), while for blue points

To more directly assess whether the same sperm sub-population was driving these correlations we designed a double labeling experiment (Figure 5C). For this experiment single stranded DNA nicks were labeled via Nick translation with FITC, while accessible chromatin regions were labeled with ATTO565-dATP (a red fluorescent dye). The presence of 5-methylcytosine containing dNTPs in the initial Nick translation reaction ensured that DNA synthesized downstream of endogenous nicks was not further cleaved by Nt.CviPII during the subsequent reaction and thus should not lead to removal signal in these regions^24^. To control for possible incomplete repair following the initial labeling reaction, we also performed a sequential Nick translation reaction, where the incorporation of ATTO565-dATP is expected to be minimal. We observed that sperm with very high levels of DNA nicking did tend to show some increased ATTO565 signal (Figure 5D red points), suggesting that these represent sperm with extensively damaged DNA that T4 DNA ligase was not able to fully repair. Nonetheless, we observed multiple populations of sperm in our double labeling experiments (Figure 5D blue points). In both total and swim-up sperm samples, we found a population possessing both high DNA accessibility and high DNA nicking signal, as well as a population with high DNA accessibility but limited DNA nicking (Figure 5D). This suggests that high sperm DNA accessibility is not always accompanied by the presence of DNA damage. We failed to detect a population of sperm with low DNA accessibility and high DNA nicking, but this may be caused by technical challenges as Nick translation may require at least some accessibility to enable sufficient incorporation of fluorescent label for detection.

## Discussion

Mammalian sperm chromatin is highly specialized for the effective and safe delivery of genetic material to the next generation. This involves packaging the DNA into an especially dense structure. Here we report that in humans the level of DNA packaging in single sperm varies dramatically within and between samples from individuals of differing reproductive outcomes, with two states being suggested by quantification of signal intensities (Figure 1C). Interestingly, the groups of infertile individuals in our cohort were separated by differences in early embryonic development, not by differences in fertilization rate. As such, our findings are among the first to demonstrate that differences in sperm chromatin packaging in infertile men may impact the likelihood of embryonic pre-implantation embryonic development.

Our results raise several interesting questions related to the generation of these varying states and their potential functional differences. Firstly, the finding that sperm selected for motility via swim-up show a general increase in DNA accessibility was unexpected. Since its first description more than 50 years ago^53^, the swim-up method has been widely used as a method for isolation of motile sperm. Previous studies have found that swim-up is an effective approach to eliminate sperm with DNA fragmentation^51,52^, which we confirmed in our Nick translation results. Swim-up and density gradient centrifugation remain the two most used sperm preparation methods for clinical ART, with studies suggesting similar reproductive outcomes from both methods following intrauterine insemination^54^ or IVF/ICSI^55^. The increase in highly accessible sperm following swim-up preparation may arise from two possible sources. The process of preparing sperm may alter chromatin, such that accessibility is increased, or alternatively sperm with more accessible DNA may be enriched within the motile fraction of total sperm. Further experiments using different sperm preparation methods will be needed to separate these two different possibilities.

A second open question relates to functional differences between NicE-view^high^ and NicE-view^low^ sperm. Overall, the frequency of NicE-view^high^ sperm in both total and swim-up samples across all groups correlates negatively with conventional semen parameters (Figure 1C). Interestingly, our participant cohort was selected such that these conventional parameters were generally within the 5^th^ percentile of the WHO fertile reference range^27^. This suggests that even within these fertile ranges further subdivision of values may contain information related to sperm quality. In natural pregnancies, the sperm concentration shows a clear relationship to time to pregnancy in low ranges (below 40-50 million/ml), but above those values increasing sperm quantity does not improve outcomes^56,57^. It is interesting to wonder if combining NicE-view measurements with conventional semen parameters would refine the predictive power of semen analysis for reproductive outcome, especially following ART. We observed that total sperm samples containing very high frequencies of NicE-view^high^ sperm most commonly occurred in the LBGR sub-group, but the number of individuals with this pattern was uncommon even among this category (only 5 of 20). Subjecting sperm samples to swim-up preparation revealed differences that were not seen in total samples in more than 30% of individuals, independent of participant category (Figure 2B). What is the difference between “Double low” and “Swim-up high” samples? Our analysis again suggests decreased sperm concentration, but also increased DNA nicking in these samples. The origin of these nicks may provide a further explanation as to the establishment of the NicE-view^high^ state that we observe in some sperm.

The chromatin state of haploid round spermatids is similar to that found in somatic nuclei, enriched for histone proteins and transcriptionally active^58^. During haploid male germ cell development (spermiogenesis) this chromatin state undergoes dramatic re-configuration, with the vast majority of histone proteins being removed and replaced by protamines, and transcription being globally silenced^59^. Topoisomerase enzymes, which maintain DNA topology, are thought to be important for this process^60^. The enzymatic activity of Class I topoisomerases generates transient single-stranded DNA breaks^61^, while Class II enzymes generate double stranded breaks to relieve DNA supercoiling^62^. These DNA breaks are usually rapidly repaired following topoisomerase reactions, but failure in this process could lead to persistent DNA damage (such as the Nick translation signal observed in some of our samples). Enzymatic activity of both classes of topoisomerases has been detected in mature human sperm^63^. Sperm in which the proper chromatin re-configuration process is aberrant may fail to fully incorporate protamines into their genomes, in addition to maintaining persistent DNA damage.

Whether the variation in chromatin state observed in mature sperm arises in the testis during spermiogenesis or develops later during epididymal transit or even in the vas deferens or after ejaculation remains unknown. The absence of a correlation between abstinence time and NicE-view^high^ frequency (Figure 2C) suggests that this state does not increase with sperm aging. Characterization of the chromatin of spermatids or testicular sperm by NicE-view would be interesting to address this question. Testicular sperm has been found to function more effectively in ART in individuals where DNA fragmentation in ejaculated sperm is elevated^64,65^. Our cohort does not possess globally increased levels of sperm DNA fragmentation as measured by TUNEL (Table 1), but if NicE-view^high^ sperm are not found in testicular sperm, it is intriguing to wonder if isolation of sperm via testicular sperm extraction could be used in cases where pre-implantation development following ART fails.

## Author Contributions

M.E.G., C.D.G. and A.H.F.M.P. conceived and planned the project. C.D.G. recruited study participants. M.F. supervised processing of human sperm samples and generation of conventional semen analyses and TUNEL assays. M.E.G. performed microscopy experiments, performed FACS isolation of CMA3 stained sperm, analyzed data and wrote the manuscript. All authors edited and approved the manuscript.

## Acknowledgements

We thank all participants for providing the samples without which this research would not be possible. We thank the staff of the RME Andrology laboratory (Universitätsspital Basel) for processing samples and performing semen analysis and TUNEL assays. We thank Steven Bourke, Laure Plantard and Laurent Gelman (FMI) for their advise and assistance in microscopy experiments. We thank Hubertus Kohler (FMI) for advice and assistance with FACS and Grigorios Fanourgakis (FMI) for the production of ZZ-tagged TN5 protein and advice on ATAC experiments. We thank the members of the Peters’ group (FMI) for useful discussion and advice on the manuscript. This work was supported by Swiss National Science Foundation Grant #189264, the Swiss Center for Applied Human Toxicology research grant no. 1 “male reproductive toxicity” (to C.D.G.) and the Novartis Research Foundation (to A.H.F.M.P.).

