## Supplementary Material for "Levels of DNA accessibility in human sperm vary across individuals with differing reproductive parameters"

### Gill et al. Supplementary Figure S1

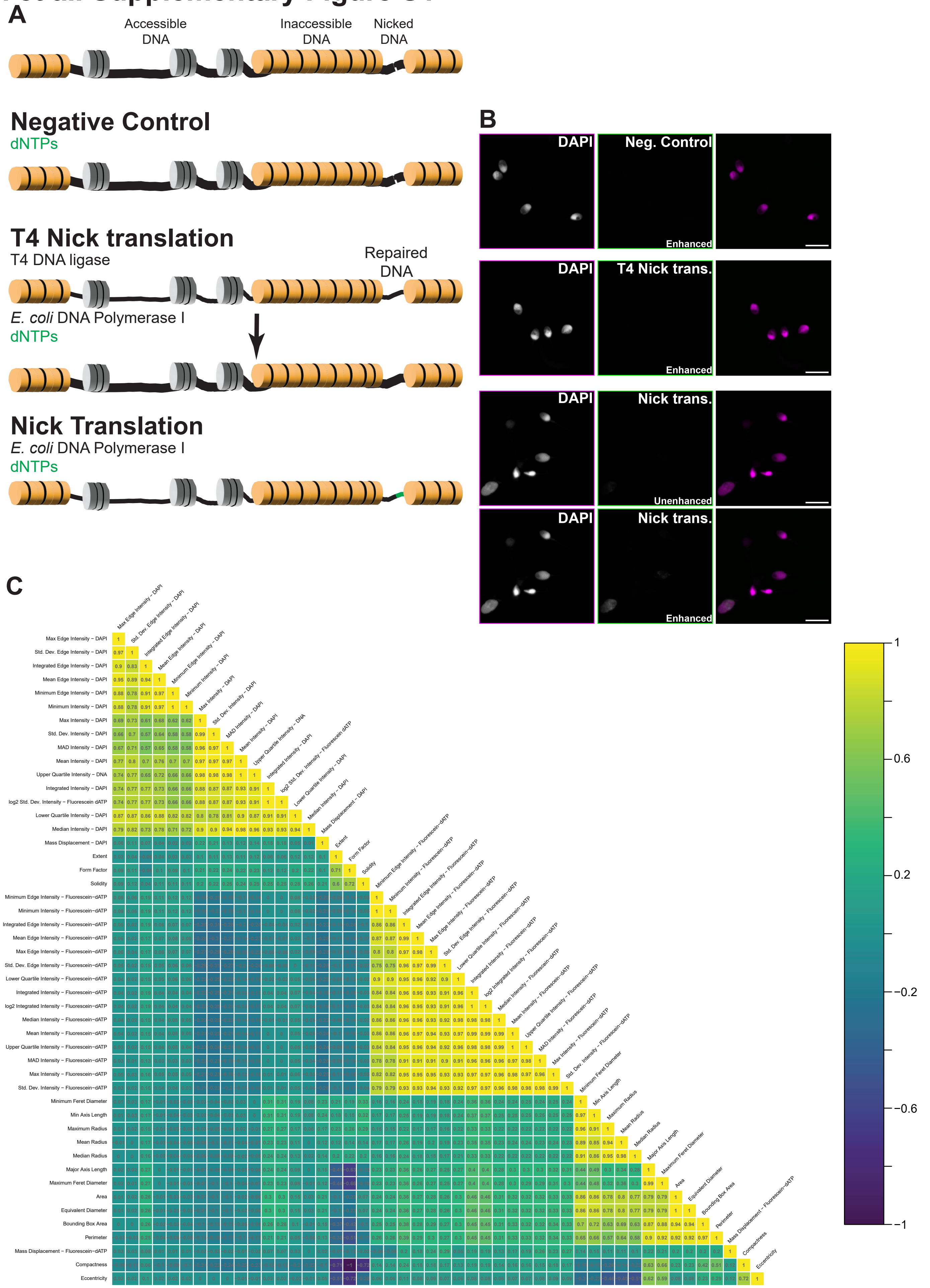


**Supplementary Figure S1 (Related to Figure 1). Control staining shows low background of NicE-view and effectiveness of T4 DNA ligase.** (A) & (B) as in Figure 1(A&B) but including two additional staining conditions: Negative control (incubated with fluorescent dNTPs and no enzymes) and T4 Nick translation (treated with T4 DNA ligase prior to Nick translation reaction). Unenhanced refers to images with no adjustment to brightness and contrast, while enhanced refers to images where brightness and contrast have been enhanced to make Nick translation signal more visible. Scale bars = 10 μm. (C) Correlation plot for all object-level imaging parameters measured for NicE-view data. Values represent Spearman's ρ.

### Gill et al. Supplementary Figure S2

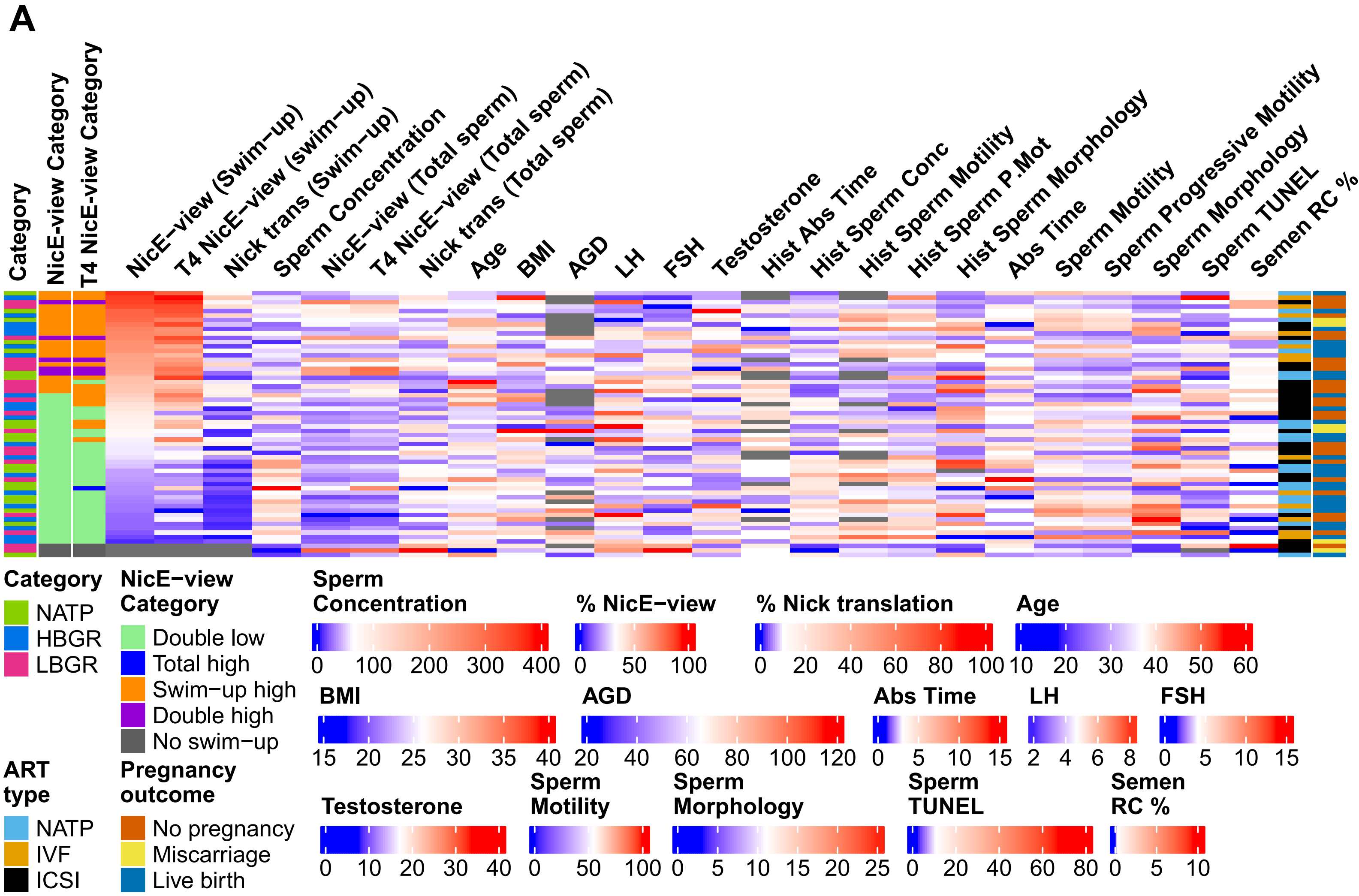


**Supplementary Figure S2 (Related to Figure 3). No correlation of NicE-view frequency with demographic or hormonal parameters in participant cohort.** Heatmap as in Figure 3C showing additional parameters related to participant cohort. Data are ordered by % NicE-view$^{high}$ sperm in swim-up sample.

### Gill et al. Supplementary Figure S3

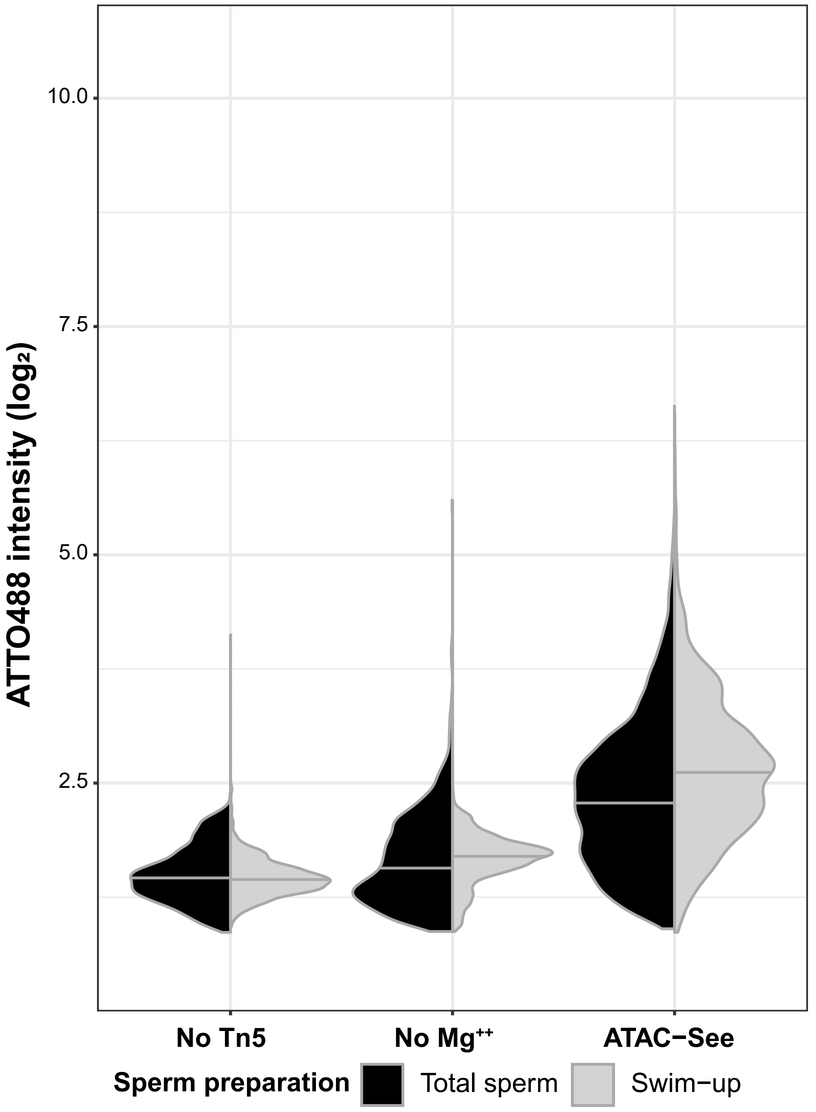


**Supplementary Figure S3 (Related to Figure 4). Tn5 enzymatically integrates labelled oligonucleotides into sperm chromatin.** (A) Split violin plot showing control reactions for ATAC-See. No Tn5 refers to incubation with labeled oligonucleotides but no enzyme; No $Mg^{++}$ refers to incubation with labeled oligonucleotides and Tn5 transposase without Magnesium (a necessary Tn5 cofactor). Data are pooled from 5 individuals.
